# With the leisure of time, kinetic proofreading can still perform reliable ligand discrimination

**DOI:** 10.1101/2024.01.11.575129

**Authors:** Fangzhou Xiao, Vahe Galstyan

## Abstract

In a recent PNAS publication, Kirby and Zilman studied kinetic proofreading for receptor signaling and argued that having more proofreading steps does not generally improve ligand discrimination when stochastic effects are taken into account. In their analysis, however, the authors made an implicit assumption about a fixed discrimination time, which is not always warranted. We show that when the discrimination time is allowed to vary in order to maintain a fixed level of signaling activity, the proofreading performance can, in fact, uniformly improve with the number of steps. We therefore call for a comprehensive study of the interplay between discrimination time, discrimination accuracy, and signaling activity, where the two seemingly contradictory results will be slices of the trade-off surface along different axes.

---

Kinetic proofreading is a canonical scheme believed to be responsible for the high ligand discrimination capacity of many biochemical processes. In a recent PNAS publication, Kirby and Zilman studied kinetic proofreading for receptor signaling, and argued that when stochasticity is taken into account, having more proofreading steps does not generally improve the discrimination of different receptor-binding ligands based on counts of signaling molecules produced [1].

In their analysis, however, the authors made an implicit assumption about a fixed discrimination time, i.e. the time to produce signaling molecules before the cell is asked to discriminate is kept the same when the number of proofreading steps N is varied. Indeed, in such a setup, the distributions of signaling molecule counts arising from cognate (lower k_off_) and noncognate (higher k_off_) ligand binding tend to overlap more with increasing N (Fig. 1a), which results in decreasing discrimination performance, as measured by the probability of false activation (Fig. 1c). This assumption, however, is not always warranted, since discrimination time could naturally change with N due to feedback [2] or signaling threshold mechanisms [3, 4].

**Figure 1:**
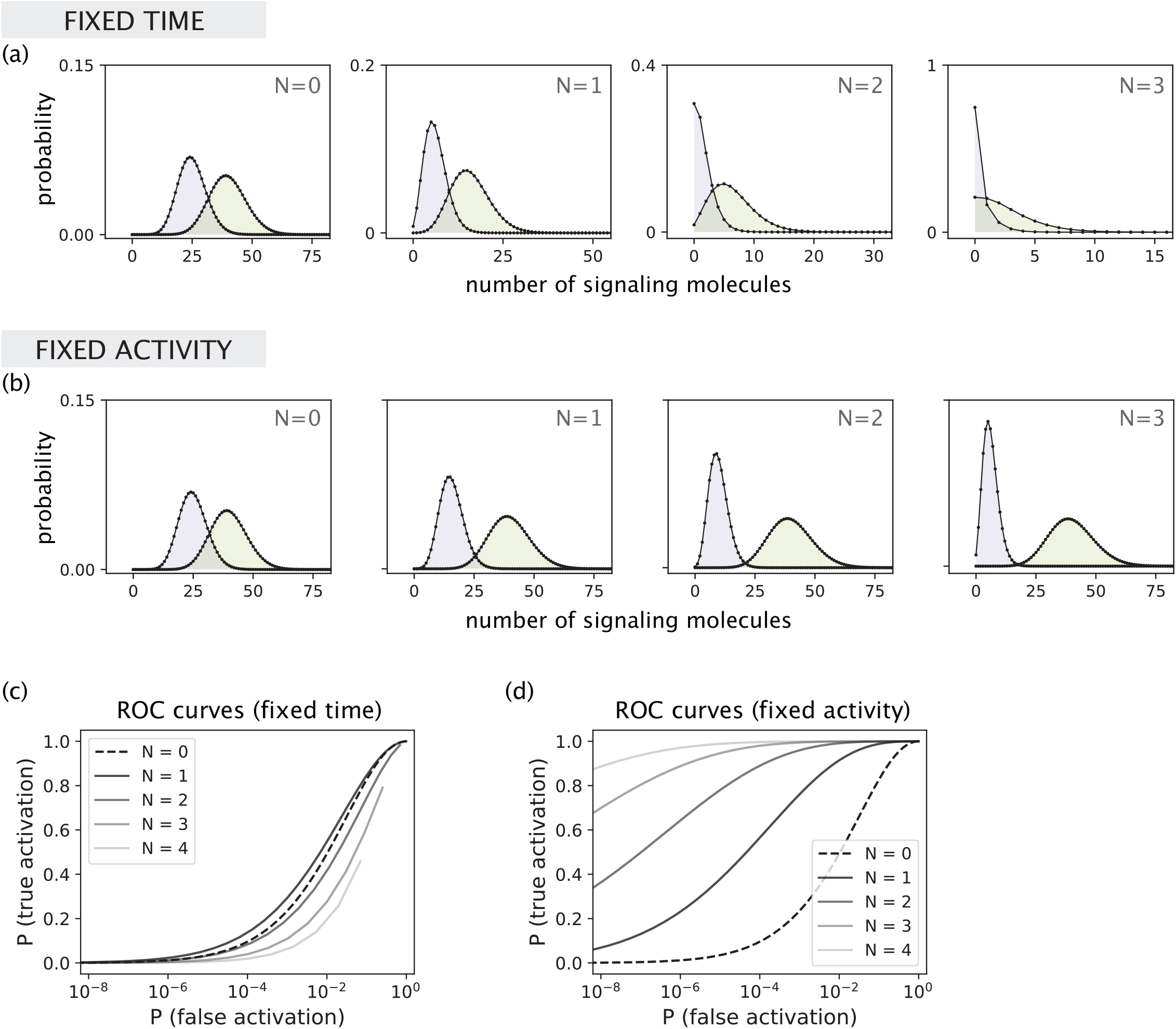
Discrimination performance can exhibit diverse responses to varying the number of proofreading steps. **(a)** Signal distributions arising from the binding of cognate (lower k_off_) and non-cognate (higher k_off_) ligands for different choices of the number of proofreading steps N in the fixed time setting. Parameter used are the same as in Fig. 2B of Kirby and Zilman: k_on_c/k_p_ = 1, k_f_/k_p_ = 1, k_p_t = 100, k_off,1_/k_p_ = 3, and k_off,2_/k_p_ = 1.5. Distributions are obtained from solving the full master equation in Eq. [11] of Kirby and Zilman (see [5] for the code). **(b)** Signal distributions for different choices of N in the fixed activity setting where discrimination time t_N_ scales with N as t_N_ = t_0_(1 + k_off,2_/k_f_)^N^ with k_p_t_0_ = 100. All other parameters are the same as in panel (a). **(c)** ROC curves for different choices of N corresponding to panel (a). The true activation probability for a given threshold n^∗^ is computed as P(n > n^∗^ | k_off,2_), and the false activation probability is computed as P(n > n^∗^ | k_off,1_). **(d)** ROC curves corresponding to the fixed activity setting in panel (b).

Here, we show that when the discrimination time is allowed to vary, the proofreading performance can, in fact, uniformly improve with the number of steps. Specifically, we consider a fixed activity setup where discrimination time increases with N to maintain the same mean level of signaling molecules produced from cognate ligand binding. This is a biologically motivated scenario where a fixed threshold of signaling activity is required for downstream decision making. While the signaling molecule count distribution corresponding to cognate ligand binding remains mostly unchanged as N is increased, the distribution corresponding to noncognate ligand binding uniformly drifts towards zero, reducing the overlap between the two distributions (Fig. 1b). As a result, discrimination performance improves with N (Fig. 1d), which is the opposite of the fixed time behavior considered by Kirby and Zilman.

While fixing signaling activity is a natural choice for thresholding-based schemes, it is arguable that fixing the discrimination time could also be biologically relevant when there is a time constraint for eliciting cellular response. Kirby and Zilman’s work therefore opens an avenue for a comprehensive study of the interplay between discrimination time, discrimination accuracy and signaling activity, where the two cases discussed here will be slices of the trade-off surface along different axes.

